# Microenvironmental Snail1 is a driver of immunosuppression in melanoma

**DOI:** 10.1101/2022.11.23.517623

**Authors:** Marta Arumi-Planas, Francisco Javier Rodriguez-Baena, Francisco Cabello-Torres, Francisco Gracia, Cristina Lopez-Blau, M. Angela Nieto, Berta Sanchez-Laorden

**Author notes:** Author for correspondence: Berta Sanchez-Laorden.

## Abstract

Melanoma is an aggressive form of skin cancer due to its high metastatic abilities and resistance to therapies. Melanoma cells reside in a heterogeneous tumour microenvironment that acts as a crucial regulator of its progression. Snail1 is an epithelial-to-mesenchymal transition transcription factor expressed during development and reactivated in pathological situations including fibrosis and cancer. In this work, we show that Snail1 is activated in the melanoma microenvironment, particularly in fibroblasts. Analysis of murine models that allow stromal Snail1 depletion and therapeutic Snail1 blockade indicate that targeting *Snail1* activation in the tumour microenvironment decreases melanoma growth and lung metastatic burden, extending mice survival. Transcriptomic analysis of melanoma-associated fibroblasts and analysis of the tumours indicate that stromal Snail1 induces melanoma growth by promoting an immunosuppressive microenvironment and pro-tumour immunity. This study unveils a novel role of Snail1 in melanoma biology and supports its potential as a therapeutic target.

## Introduction

Melanoma is the most aggressive form of skin cancer. If found early, it can be surgically resected, but melanoma is extremely metastatic and very resistant to treatments when disseminated to other organs. Even though in recent years the landscape of melanoma treatment has greatly improved with the use of more effective targeted therapies and immunotherapies, not all patients respond to these treatments and many of the patients who respond develop resistance after a relatively short period of disease control^1^. Importantly, melanoma progression and how it responds to treatments is strongly influenced by the tumour microenvironment (TME)^2^.

Epithelial to mesenchymal transition (EMT) is a developmental process that can be triggered in pathological conditions including fibrosis and cancer. Epithelial cells undergo EMT acquiring the capacity to move and disseminate^3,4^. EMT endows cancer cells with invasive and migratory capabilities as the tumour progresses^5,6^. The main inducers of the EMT are transcription factors (TFs) of the Snail, Twist and Zeb families. EMT-TFs coordinate the downregulation of epithelial genes and the induction of mesenchymal ones^3,7^. EMT-TFs play an important role in the development and dissemination of epithelial-derived carcinomas, particularly when they are expressed in tumour cells^3,8,9^ but also when its expression is associated with stromal cells, particularly cancer-associated fibroblasts (CAFs)^10–14^. CAFs, central components of the tumour stroma, are a complex and heterogeneous population of myofibroblasts whose activity associates with tumour aggressiveness. CAFs coordinate a wide array of functions including matrix remodelling, angiogenesis, and tumour-promoting immune evasion^15,16^.

Reprogramming in the expression of different EMT-TFs, including Zeb1 and 2 and Twist or Snail2 in melanoma cells is associated with tumour progression^17–21^. In addition, previous studies have assessed the impact of Snail1-induced EMT in melanoma cells^22–24^. However, whether Snail1 expression in the TME regulates melanoma biology has not been investigated. In this study, we use different mouse models to unveil a novel immunoregulatory role of Snail1 reactivation in the melanoma microenvironment. We show that Snail1 expression in fibroblasts regulates fibrillary acid protein (Fap) expression and promotes immunosuppression. Consistent with the latter, Snail1 targeting significantly decreases tumour and metastatic burden, increasing mice survival. We also show that the effects driven by microenvironmental Snail1 targeting are associated with an increase in anti-tumour immune responses. Altogether, this indicates that stromal Snail1 has a crucial role in shaping the melanoma microenvironment to drive tumour progression.

## Results

### Snail1 reactivation in the tumour microenvironment promotes melanoma growth

Snail1 expression has been previously found in epithelial and stromal cells in carcinomas^11,25^. To characterise the expression of Snail1 in melanoma, and to distinguish tumour cells from the cells in the TME, we generated a melanoma reporter mouse model by crossing the inducible BRAF-driven mouse melanoma model BRaf^CA^,Pten^loxP^,Tyr::CreERT2 (BRAF^V600E^/Pten^loxP^)^26^ with Rosa-LSL-tdTomato mice (tdTomato). Tamoxifen treatment of these mice results in melanoma development with a short latency^26^ and the expression of the Tomato fluorescent protein in melanocytes and melanoma cells. Analysis of this model showed SNAI1 expression restricted to tdTomato-negative cells in the tumours (Fig. 1a, b) indicating that *Snail1* is reactivated in the melanoma microenvironment but not in the melanoma cells. To specifically target the stroma, we next generated a syngeneic melanoma model by injecting murine Braf^V600E^-5555 cells^27,28^ in UBC-Cre-ERT2 mice^29^ crossed with tdTomato mice (Fig. 1c). In this model, tamoxifen treatment promotes the ubiquitous expression of the Tomato fluorescent protein in the mouse, allowing to trace the red labelled stromal cells in the allografts. Analysis of the tumours confirmed SNAI1 expression in the recombined cells from the melanoma microenvironment that was absent in normal skin (Fig. 1d). We extended our analyses to additional oncogenic BRAF and BRAF^wt^/NRAS^wt^ melanoma syngeneic models and confirmed SNAI1 reactivation in the stroma of these tumours (Supplementary Fig. 1). Next, we wanted to assess the contribution of microenvironmental Snail1 (Snail^ME^) to melanoma growth. For this, UBC-Cre-ERT2-tdTomato mice were bred with *Snai1^fl/fl^* mice^30^ to prevent Snail1 reactivation in the tumour stroma. Melanomas were established by subcutaneous injection of Braf^V600E^-5555 cells in UBC-Cre-ERT2-tdTomato and UBC-Cre-ERT2-tdTomato-Snai1^*fl/fl*^ (referred as Snail1^ME^-WT and Snail1^ME^-KO, respectively) (Fig. 1c). When the tumours were already established, animals were treated with tamoxifen to block stromal Snail1 expression and melanoma growth was monitored (Fig. 1e). We confirmed that recombined stromal cells from Snail1^ME^-KO mice lack SNAI1 expression (Fig. 1f). Importantly, melanoma growth was blocked and significantly reduced in Snail1^ME^-KO compared to Snail1^ME^-WT mice (Fig. 1g, h). In line with these results, we observed a decrease in the proliferation of melanoma cells from Snail1^ME^-KO tumours (Fig.1i, j) and a significant increase in apoptotic melanoma cells as indicated by cleaved-Caspase 3 (Fig 1k, l). These results show that Snail1 is expressed in the melanoma microenvironment where it is necessary for melanoma growth.

**Figure 1.**
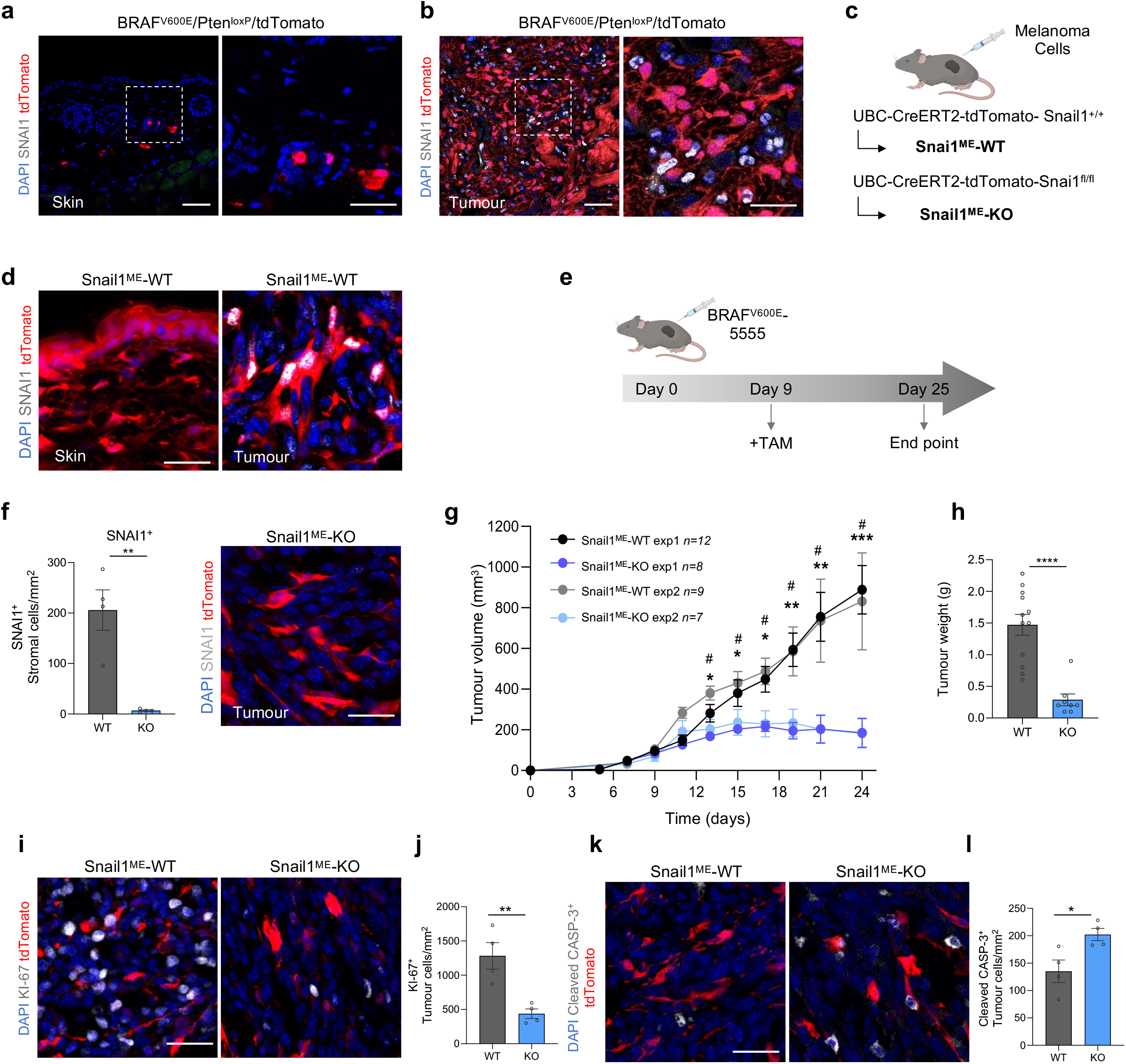
Snail1 is expressed in the melanoma microenvironment and its ablation reduces tumour growth and promotes apoptosis. **(a,b)** Representative images of immunolabelling for SNAI1 (white) in control skin and tumours from BRaf^CA^,Pten^loxP^,tdTomato,Tyr::CreERT2 (BRAF^V600E^/Pten^loxP^/tdTomato) mice. Melanoma cells are labelled in red (tdTomato). **(c)** Mouse models generated to investigate the impact of Snail1 on the melanoma microenvironment. **(d)** Representative images of immunolabelling for SNAI1 (white) in control skin (left panel) and Braf^V600E^-5555 tumours (right panel) from Snail1^ME^-WT mice. Stromal cells are labelled in red (tdTomato). **(e)** Experimental set-up of the *in vivo* strategy design to study the contribution of Snail1 to melanoma progression. Created with BioRender.com. **(f)** Quantification of SNAI1^+^ stromal cells (n=4 per condition) (left panel) and representative image of immunolabelling for SNAI1 (white) in Snail1^ME^-KO tumours upon tamoxifen administration (right panel). Stromal cells are labelled in red (tdTomato). **(g)** Braf^V600E^-5555 tumour growth was assessed in two independent experiments combined in this graph (exp1 n=12 Snail1^ME^-WT and n=8 Snail1^ME^-KO; exp2 n=9 Snail1^ME^-WT and n=7 Snail1^ME^-KO). **(h)** Final weight after collection of tumours from Snail1^ME^-WT (WT) and Snail1^ME^-KO (KO) mice (n=12 WT; n=8 KO). **(i)** Representative images of immunolabelling for KI-67(white) in tumours from Snail1^ME^-WT and Snail1^ME^-KO mice. Stromal cells are labelled in red (tdTomato). **(j)** Quantification of KI-67 (white) tumour nuclei-positive cells in images from (i) (n=4). **(k)** Representative images of immunolabelling for Cleaved-Casp3 (white) in tumours from Snail1^ME^-WT and Snail1^ME^-KO mice. Stromal cells are labelled in red (tdTomato). **(l)** Quantification of images from (k) (n=4). Data are represented by Mean±SEM and statistically significant differences are tested by unpaired two-tailed Student t-test. Each dot represents one animal (*=p<0.05, **=p<0.01, ***p<0,001, ****p<0,0001 and #=p<0.05 for experiment 2). WT=Snail1^ME^-WT and KO= Snail1^ME^-KO. Scale bars: 50μm and 25μm for higher magnification pictures.

### Snail1 reactivation in melanoma-associated fibroblasts decreases anti-tumour immunity

Expression of Snail1 and other EMT-TFs have been reported in macrophages and CAFs from epithelial-derived tumours^10–14^. To investigate Snail1 expression in these cell populations in melanoma, we first analysed Braf^V600E^-5555 tumours grown subcutaneously in Cx3cr1CreERT2-YFP reporter mice. These mice constitutively express YFP in the myeloid lineage including monocytes and macrophages^31,32^. We did not detect SNAI1 on myeloid cells in our tumours (Fig. 2a) or in additional immune populations as assessed by CD45 staining (Fig. 2b, Supplementary Fig. 2). On the contrary, we detected SNAI1 expression in melanoma-associated fibroblasts, as indicated by double tdTomato-PDGFRα positive staining (Fig. 2c). SNAI1 positive expression in melanoma-associated fibroblasts was further confirmed in the melanoma transgenic BRAF^V600E^/Pten^loxP^/tdTomato model (Fig. 2c).

**Figure 2.**
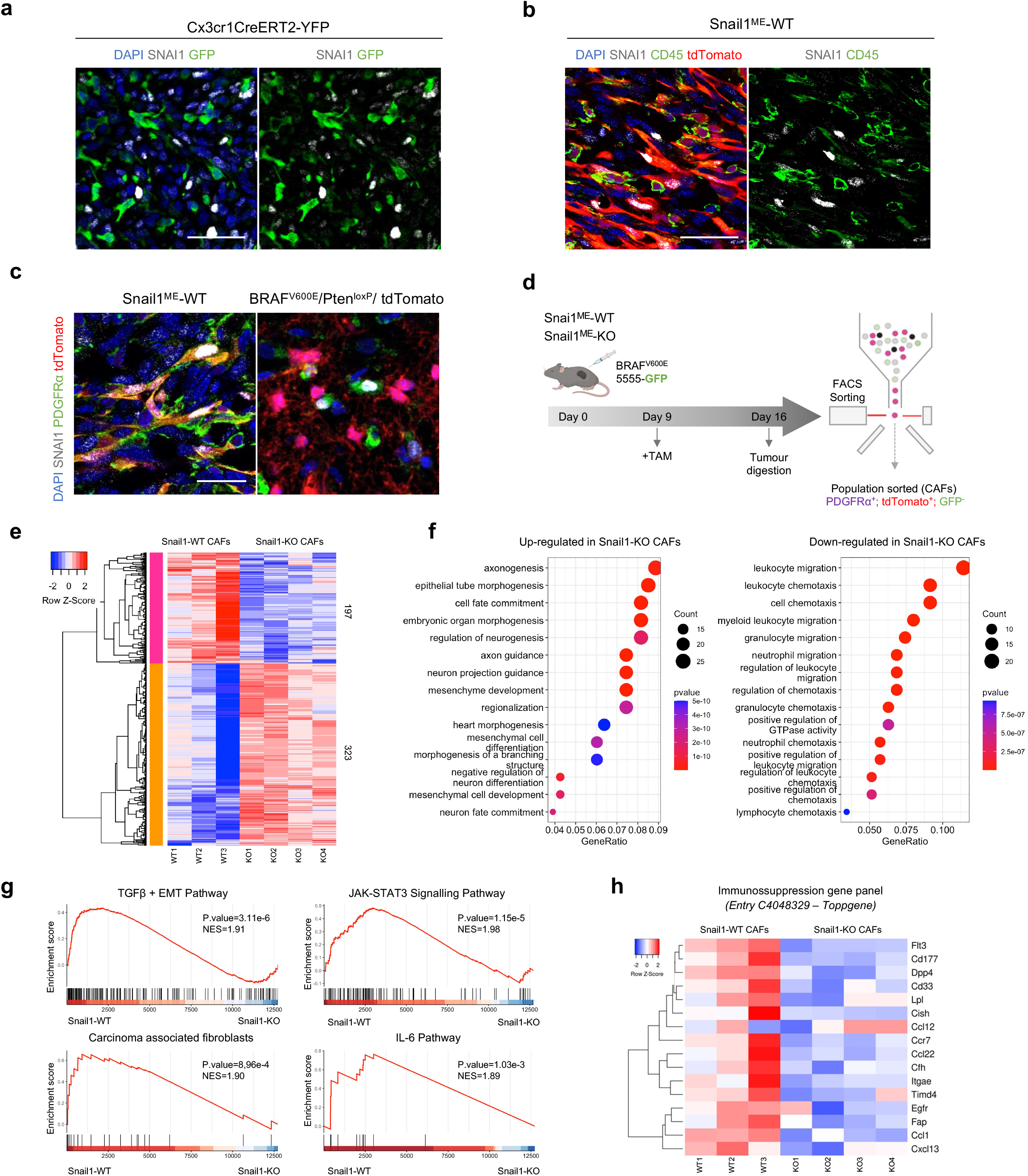
Snail1 expression in PDGFRα^+^-CAFs is associated with fibroblast activation and immunosuppression signatures. **(a)** Representative images of immunolabelling for SNAI1 (white) and myeloid cells (green) in a section of a Braf^V600E^-5555 melanoma grown in Cxcr1CreERT2-YFP mice. **(b)** Representative images of immunolabelling for SNAI1 (white) and CD45 (green) in melanomas from Snail1^ME^-WT mice. Stromal cells are labelled in red (tdTomato). **(c)** Representative images of immunolabelling for SNAI1 (white) and PDGFRα (green) in Braf^V600E^-5555 tumours from Snail1^ME^-WT mice. Stromal cells are labelled in red (tdTomato) (left panel) and in BRAF^V600E^/Pten^loxP^/tdTomato melanomas where melanoma cells are labelled in red (tdTomato) (right panel). Scale bar: 25μm. **(d)** Schematic illustration of the strategy followed to isolate fibroblasts from Braf^V600E^-5555 melanomas in Snail1^ME^-WT and Snail1^ME^-KO mice. Created with BioRender.com. **(e)** RNA seq heatmap of differentially expressed genes (DEGs). The scale bar corresponds to row Z score in a −2 to 2 relationship. Filtered and normalised count per million data from the DEGs has been plotted to compare Snail1-WT and Snail1-KO CAFs. Columns represent the different samples. Each sample is a pool of three different animals with the same genotype WT n= 3, KO n= 4. **(f)** Representation of gene ontology enrichment analysis of the 15 top GO terms as ranked by various gene set testing methods. The dot plot size and colour represent the relative number and relevance of the genes in the set, respectively. **(g)** Gene set enrichment analysis (GSEA) of DEGs genes (log2 ratio-ranked) shows enrichment of TGFβ + EMT, JAK-STAT3 and IL-6 pathway signatures and enrichment of Carcinoma-associated fibroblasts signature in Snail1-WT CAFs. NES (normalised enrichment score) and p-value scores are shown. **(h)** Panel showing expression of genes associated with immunosuppression from entry C4048329 in the Toppgene and DisGeNet databases.

To determine the mechanisms implicated in Snail1 contribution to melanoma growth, we isolated tdTomato^+^PDGFRα^+^ cells from Braf^V600E^-5555 tumours grown in Snail1^ME^-WT and Snail1^ME^-KO mice after tamoxifen treatment and performed RNA sequencing (Fig. 2d, Supplementary Fig. 3a). We corroborated that isolated cells were positive for PDGFRα, SNAI1, and αSMA, an additional CAF marker (Supplementary Fig. 3b), and confirmed *Snai1* downregulation in tdTomato^+^PDGFRα^+^ cells from Snail1^ME^-KO mice (Supplementary Fig. 3c). Among the 520 differentially expressed genes (DEGs) detected upon Snail1 targeting, 323 were upregulated and 197 downregulated (Fig. 2e). In agreement with the role of Snail1 in embryonic development, gene ontology (GO) analysis of the upregulated genes showed an enrichment in biological processes associated with morphogenesis and differentiation (Fig. 2f)^33,34^. On the contrary, 11 out of the 15 most enriched biological processes in the downregulated genes were associated with the immune system (Fig. 2f). In addition, gene set enrichment analysis (GSEA)^35^ showed that melanoma-associated fibroblasts were enriched in signatures related to TGF-β signalling and fibroblast activation in carcinomas. Interestingly, this correlation was decreased upon Snail1 targeting (Fig. 2g). We also found that several of the downregulated genes including *Ccl1, Ccl22, Cxcl13* or *Ccr7* were associated with immunosuppression and decreased anti-tumour immunity^36–40^ (Supplementary Fig. 3d, e). Further, additional GO and GSEA analyses confirmed a significant decrease in processes and genes associated with immunosuppression and pro-inflammatory pathways in Snail1 depleted melanoma-associated fibroblasts^41^ (Fig. 2g, h, Supplementary Fig. 3e).

Our data suggest that the anti-tumour effects observed upon Snail1 targeting in the melanoma microenvironment may be related to CAFs immunoregulatory functions. To test this hypothesis, we characterised the immune infiltration in Braf^V600E^-5555 melanomas upon Snail1^ME^ depletion (Fig. 3a). We observed that compatible with the impaired growth of melanomas, the percentage of tumour infiltrating cytotoxic T cells (CD8^+^) was elevated in tumours from Snail1^ME^-KO compared to Snail1^ME^-WT mice (Fig. 3b). In addition, significantly fewer regulatory T cells (FOXP3^+^) were found in Snail1^ME^-KO tumours (Fig. 3c). Further analyses show an increase in B cells and Natural killer (NK) cells in tumours from Snail1^ME^-KO mice, while the number of dendritic cells and myeloid cells remained constant (Fig. 3d, e). However, we detected upregulated expression of Arginase 1 *(Arg1)* (Fig. 3f), a marker associated with M2-like macrophages with immunosuppressive and pro-tumorigenic functions, in melanomas from Snail1^ME^-WT compared to Snail1^ME^-KO mice. Altogether, these data indicate that Snail1^ME^ expression blocks anti-tumour immune responses.

**Figure 3.**
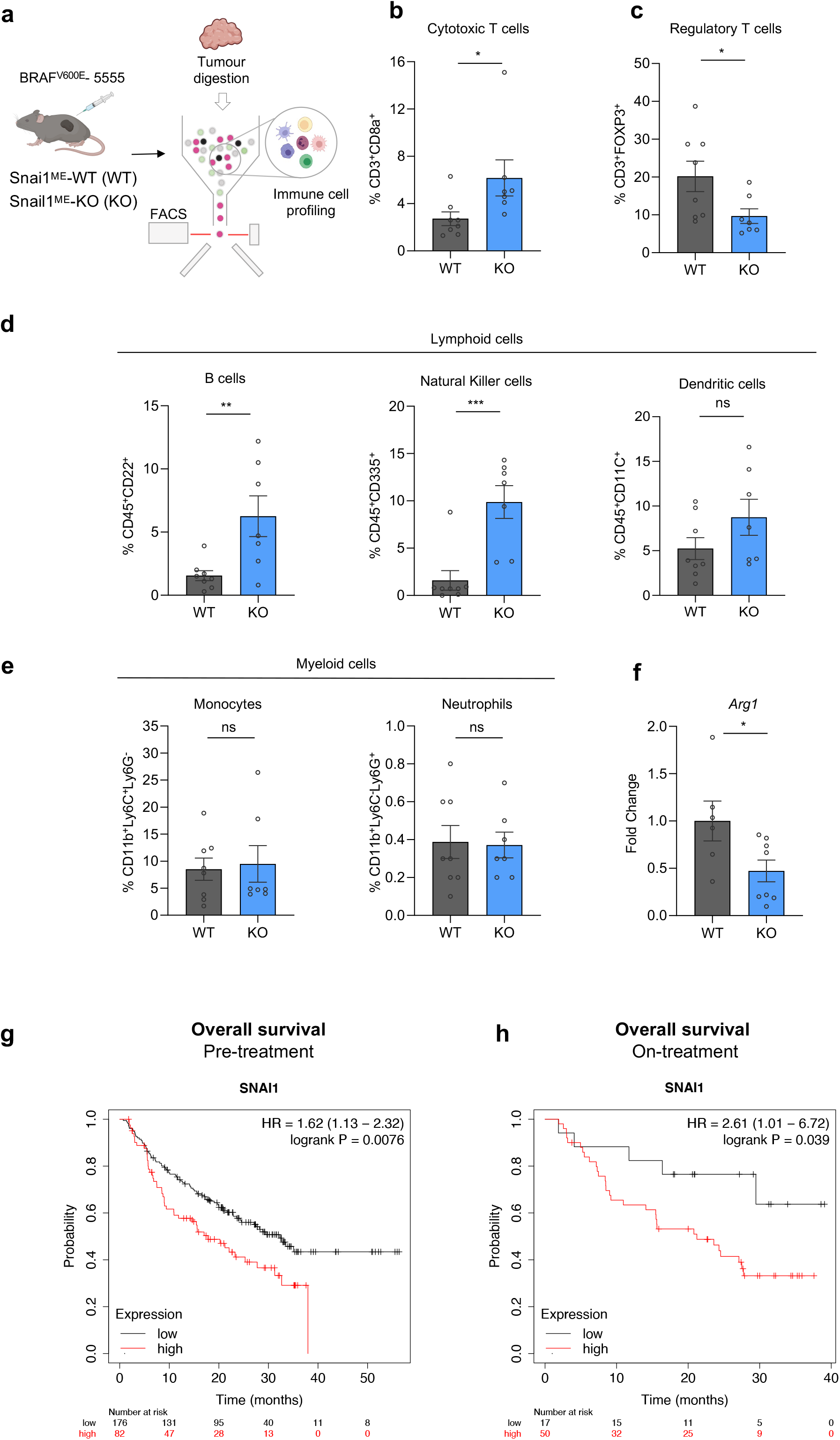
Snail1^ME^ targeting induces an anti-tumorigenic immune response in melanoma. **(a)** Schematic representation of the strategy used to perform immune cell profiling by flow cytometry analysis of Braf^V600E^-5555 melanomas in Snail1^ME^-WT (WT) and Snail1^ME^-KO (KO) mice. Created with BioRender.com. **(b, c)** Graphs showing percentages of Cytotoxic T cells (CD3^+^CD8a^+^) and Regulatory T cells (CD3^+^FOXP3^+^) in tumours from (a) (n=8 WT and n=7 KO). **(d)** Graphs showing percentages of lymphoid cells from (a); B cells (CD45^+^CD22^+^,), natural killer cells (CD45^+^CD335^+^,) and dendritic cells (CD45^+^CD11c^+^,), are represented (n=8 WT and n=7 KO). **(e)** Graphs showing percentages of myeloid cells in tumours from (a); monocytes (CD11b^+^Ly6C^+^Ly6G^-^) and neutrophils (CD11b^+^Ly6C^-^Ly6G^+^) **(f)** *Arg1* mRNA levels detected by RT-qPCR in tumour samples from (a) (n=5 WT and n=8 KO). Data are represented by Mean±SEM and statistically significant differences are tested by unpaired two-tailed Student t-test. Each dot represents one animal (ns=no significant, *=p<0.05, **=p<0.01, ***p<0,001). **(g, h)** Prognostic value of Snail1 expression in response to anti-PD-1 therapy. Survival curves plotted for melanoma patients. *Snai1* expression assessed before anti-PD-1 therapy (n=258) or on treatment (n=67). Data was analysed using Kaplan-Meier Plotter. Patients with Snail1 expression above the median are indicated in red line, and patients with expressions below the median in black line. HR, hazard ratio.

Interestingly, given our results indicating that Snail1 promotes immunosuppression in our models, and the association of immunosuppression with resistance to immunotherapy^42^, we sought to investigate the correlation of Snail1 levels with clinical outcomes in patients treated with immune-checkpoint inhibitors. Analysis from transcriptomic datasets^43^ using Kaplan-Meier Plotter^44^ revealed that high Snail1 expression before or on-treatment with anti-programmed death-1 (anti-PD-1) correlated with a lower overall survival in melanoma patients (Fig. 3g, h).

### Snail1 expression in fibroblasts activates Fap

Recent studies using syngeneic carcinoma models indicate that CAFs expressing fibroblast activation protein (FAP) are responsible for immune-evasion associated with a pro-tumorigenic TME^45–50^. Given our results showing that Snail1^ME^ in melanoma promotes immunosuppression and that its silencing blocks tumour growth, we next investigated the potential relationship between Snail1 and Fap in fibroblasts. We first analysed *Fap* expression in our models. Analysis of our RNAseq data showed a decrease in *Fap* expression in melanoma-associated fibroblast upon Snail1 depletion (Supplementary Fig. 3d). Further, we confirmed lower levels of *Fap* mRNA in tumours from Snail1^ME^-KO when compared to Snail1^ME^-WT mice (Fig. 4a). In line with this, *Fap* was downregulated in NIH3T3 fibroblasts after silencing *Snai1* expression with a siRNA (Fig. 4b), and upregulated upon TGFβ treatment or *Snai1* overexpression (Fig. 4c, d), indicating that Snail1 could be regulating Fap expression. To assess whether SNAI1 could directly bind to regulatory regions of the *Fap* promoter, we performed chromatin immunoprecipitation (ChiP) assay in *Snail1*-overexpressing NIH3T3 cells. For this, we looked for consensus Snail1 E-boxes^51^ (CANNTG) within the mouse *FAP* promoter using the SnapGene® software. Both murine and human *FAP* promoters contain multiple Snail1 E-boxes near their transcription start site. We considered regions with 2 or more E-boxes as predicted SNAI1 binding sites (BS), and we found several within the mouse *Fap* promoter (BS1, BS2, BS3, BS4 and BS5) (Fig. 4e). Chromatin immunoprecipitation analysis confirmed that BS1, BS2 and BS3 were highly enriched in SNAI1 binding as compared with IgG control in the NIH3T3 cell line (Fig. 4f). All this together indicates that SNAI1 could directly regulate *Fap* transcription in fibroblasts.

**Figure 4.**
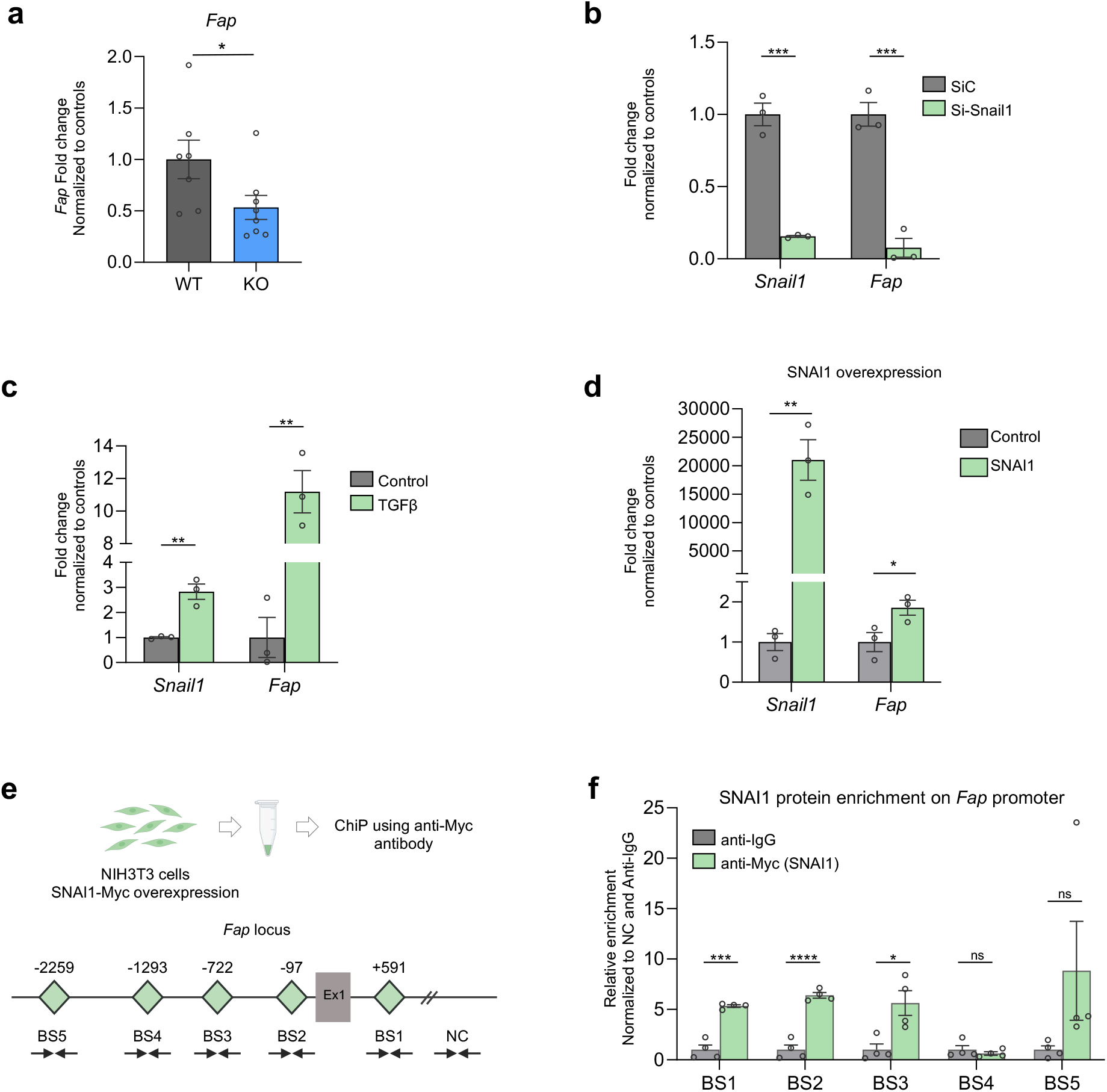
Fap is a direct target of Snail1 in fibroblasts. (a) *Fap* mRNA levels detected by RT-qPCR in Braf^V600E^-5555 tumours from Snail1ME-WT (WT) and Snail1^ME^-KO (KO) mice (n=7 WT and n=8 KO). **(b)** *Snai1* and *Fap* mRNA levels detected by RT-qPCR upon Snail1 silencing using a siRNA in TGFβ treated NIH3T3 cells. Transfected cells were collected 48h after transfection (n=3). **(c)** *Snail* and *Fap* mRNA levels increase detected by RT-qPCR upon TGFβ treatment in NIH3T3 cells. Cells were collected 48 h after TGFβ treatment (n=3). **(d)** *Snai1* and *Fap* mRNA levels increase detected by RT-qPCR after SNAI1 transfection in NIH3T3 cells (n=3). **(e)** SNAI1 enrichment on the *FAP* promoter shown by ChiP assay in NIH3T3 cells, using an anti-Myc antibody (for SNAI1-Myc overexpression). Schematic representation of the mouse *FAP* locus is shown. SNAI1 potential binding sites (E-boxes; CANNTG) on the *FAP* promoter are represented as green diamonds (BS1: +591bp, BS2: −97bp, BS3: −722bp, BS4: −1293bp, BS5 −2259bp). An intergenic region without SNAI1 binding sites was used as a negative control (NC). Ex1: SNAI1 exon 1. **(f)** Relative enrichment of SNAI1 binding to the five potential sites, normalised to the NC region and the anti-IgG controls (n=4). Data are represented by Mean±SEM and statistically significant differences are tested by unpaired two-tailed Student t-test. Each dot represents one animal (panel a) or independent experiments (panel c-f) (ns=no significant, *=p<0.05, **=p<0.01, ***p<0,001, ****p<0,0001).

### Snail1^ME^ targeting reduces metastatic burden and increases mice survival

Our results show that targeting Snail1^ME^ blocks subcutaneous melanoma growth. We next wanted to address whether Snail1 regulates the metastatic niche microenvironment. For this, we injected luciferase-expressing Braf^V600E^-5555 melanoma cells in the tail vein of Snail1^ME^-WT and Snail1^ME^-KO mice. We confirmed that SNAI1 expression was absent in control lungs and was reactivated in the metastatic microenvironment in Snail1^ME^-WT mice (Fig. 5a) and blocked in metastases from Snail1^ME^-KO mice (Fig. 5a, b). We also observed SNAI1 reactivation in PDGFRα^+^-CAFs from melanoma lung metastases in Snail1^ME^-WT mice that was confirmed in BRAF^V600E^/Pten^loxP^/tdTomato melanomas (Fig. 5c, Supplementary Fig. 4). When metastases were detected by bioluminescence in an IVIS *in vivo* imaging system, animals were treated with tamoxifen and metastases growth was monitored (Fig. 5d, e). Histological analysis of the lungs (Fig. 5f) showed a significant decrease in metastatic burden (−82,9%), metastases number (−47,6%) and size (−77,5%) in Snail1^ME^-KO compared to Snail1^ME^-WT mice (Fig. 5g). Further, we also investigated whether blocking Snail1^ME^ activation could improve mice survival. Kaplan-Meier analysis showed an almost 30% increase in the survival of Snail1^ME^-KO mice, compared to Snail1^ME^-WT, assessed by long-rank test (X^2^=6.92, p<0.01) (Fig. 5h). Importantly, as in the subcutaneous tumours, these anti-tumourigenic effects were associated with a decrease in proliferation and an increase in apoptosis in the melanoma cells in Snail1^ME^-KO metastases (Fig. 5i, j). We then analysed the immune infiltrate in the lungs and confirmed that metastases from Snail1^ME^-KO mice had an increased number of cytotoxic T cells (CD8^+^) and a lower infiltration of regulatory T cells (FOXP3^+^) compared to Snail1^ME^-WT metastases (Fig. 5k). Gene expression analysis also showed a decrease in *Fap* mRNA levels in lung metastases from the Snail1^ME^-KO mice (Fig. 5l). The data described above indicate that genetic blockade of Snail1 activation in the TME decreases metastases growth and in line with our previous results, this is associated with a less immunosuppressive environment. We had previously shown that Snail1 targeting by injection of antisense oligonucleotides could constitute a good therapeutic strategy in renal fibrosis^52^. To investigate whether this was also the case in melanoma, we used a similar approach and injected a VIVO-morpholino (VI-MO) that targets a splicing site in the *Snail1* mRNA (Snail1-MO1)^54^ into the tail vein of C57BL/6 mice with established Braf^V600E^-5555 lung metastases (Fig. 6a). Once lung metastases were detected by bioluminescence, the mice were treated with VI-MOs and the signal was monitored by IVIS (Fig. 6b). Histological analysis of the lungs (Fig. 6c) showed a decrease in the weight (−56,7%), metastatic burden (−55,9%) and number of metastases (−37,4%) in the Snail1-MO as compared to Control-MO treated mice (Fig. 6d). We confirmed the efficacy of the morpholino in blocking *S*nail1 expression in the lung metastases (Fig. 6e, f) that was accompanied by a decrease in *Fap* levels (Fig. 6f). Further, metastases from mice treated with Snail1-MO1 had increased *CD8* compared to Control-MO treated mice (Fig. 6g). Thus, as observed in our Snail1^ME^-KO mice, Snail1 systemic inhibition was associated with an anti-tumour immune response and decreased melanoma metastatic burden.

**Figure 5.**
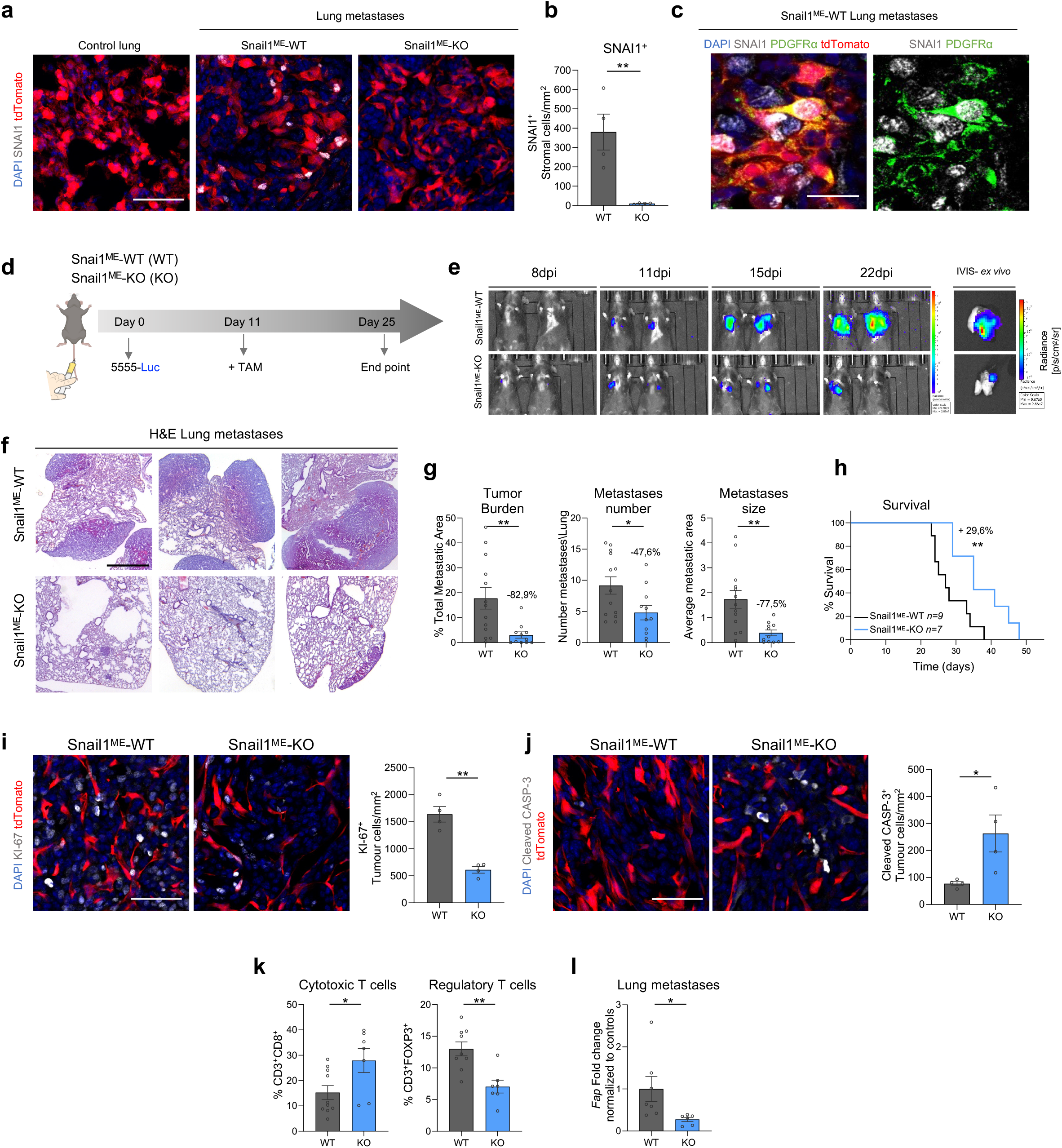
Microenvironmental Snail1 depletion reduces melanoma metastatic burden and improves mice survival. **(a)** Representative images of immunolabelling for SNAI1 in control lung tissue and Braf^V600E^-5555 lung metastases from Snail1^ME^-WT and Snail1^ME^-KO mice. Scale bar: 50 μm. **(b)** SNAI1 quantification after TAM administration in lung metastases from (a). **(c)** Representative images of double immunolabelling of SNAI1 (white) and PDGFRα (green) in Braf^V600E^-5555 lung metastases from Snail1^ME^-WT. Scale bar: 25 μm. **(d)** Experimental set-up of the *in vivo* strategy design to study the contribution of Snail1 to lung metastases progression in Snail1^ME^-WT (WT) and Snail1^ME^-KO (KO) mice. Created with BioRender.com. **(e)** Bioluminescent signal in mice from (d). The BLI scale is represented next to each panel. Units: p/s/cm2/sr (n= 13 WT and n= 11 KO). **(f)** Representative H&E-stained lung sections 25 days post-injection. Scale bar: 2mm. **(g)** Tumour burden, number of metastases and metastases size, quantified in lungs from (d) (n= 13 WT and n= 11 KO) **(h)** Overall survival of Snail1^ME^-WT and Snail1^ME^-KO mice with melanoma lung metastases after Snail1-silencing compared to controls (n= 9 WT and n= 7 KO). **(i)** Representative images of immunolabelling for KI-67 and quantification (n=4 per condition) in melanoma lung metastases from (d). Scale bar: 50 μm. **(j)** Representative images of immunolabelling for Cleaved-Caspase3 and quantification (n=4 per condition) in lung metastases from (d). Scale bar: 50 μm **(k)** Graphs showing percentages of Cytotoxic T cells (CD3^+^CD8a^+^) and Regulatory T cells (CD3^+^FOXP3^+^) in lungs from (d) (n=10 WT and n=7 KO) assessed by flow cytometry. **(l)** *Fap* mRNA levels assessed by RT-qPCR for in Snail1^ME^-WT (WT) and Snail1^ME^-KO (KO) lung metastases. Data are represented by Mean±SEM and statistically significant differences are tested by unpaired two-tailed Student t-test. Each dot represents one animal (*=p<0.05 and **=p<0.01).

**Figure 6.**
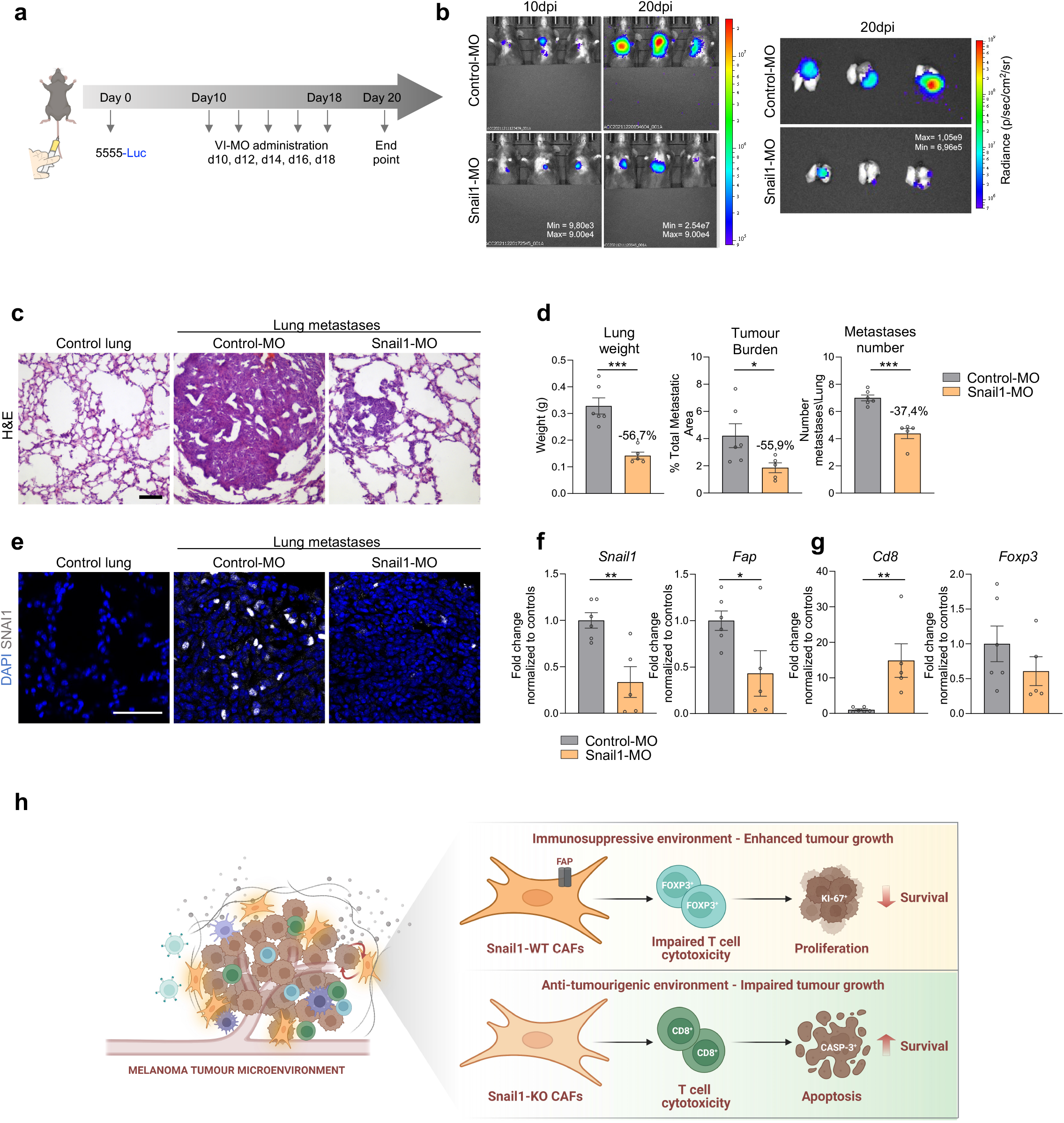
Snail1 systemic targeting significantly reduces melanoma lung metastases in mice. **(a)** Scheme of the experimental approach. Nine days after tail vein injections, C57BL/6 mice were injected with vivo-morpholino (VI-MO) control (Control-MO) or Snail1 morpholino (Snail1-MO) every other day. Created with BioRender.com. **(b)** Bioluminescent signal in mice from (a). The BLI scale is represented in each panel. Units: p/s/cm2/sr. **(c)** Representative H&E-stained lung sections after VI-MO treatment. Scale: 100μm. **(d)** Final lung weight, tumour burden and number of metastases from mice in (a) were quantified at the end of the experiment. **(e)** Representative images of immunolabelling for SNAI1 in lung sections after VI-MOs treatment. Scale bar: 50 μm. **(f,g)** *Snail1, Fap, CD8a* and *Foxp3* mRNA expression assessed by RT-qPCR lung metastases from mice treated with Snail1-MO (n=6 Control-MO and n=5 Snail1-MO). Data are normalised to samples treated with Control-MO. Data are represented by Mean±SEM and statistically significant differences are tested by unpaired two-tailed Student t-test. Each dot represents one animal (ns=no significant, *=p<0.05, **=p<0.01, ***p<0,001). **(h)** Summary illustration of the main conclusions from this study. In brief, Fap expression is regulated by Snail1 in fibroblasts, in association to an immunosuppressive phenotype in the melanoma microenvironment driven in part by cytotoxic T cell dysfunction and regulatory T cell enrichment, promoting melanoma growth. In the absence of Snail1 in the TME, Fap expression is downregulated in CAFs and a shift to an antitumour immune response occurs. Those changes are directly associated with increased melanoma cell apoptosis, smaller tumour volume, reduced metastatic burden and increased mice survival. Created with BioRender.com.

## Discussion

Modulation of the immune response in the TME plays a major role in the clinical response to treatments^53^. In this study, we have identified stromal Snail1 as a driver of melanoma growth by promoting an immunosuppressive TME. Moreover, Snail1 targeting is enough to reduce melanoma metastatic burden and increase mice survival (Fig. 6h).

Snail1 is an essential TF during embryonic development whereas it is mostly absent in healthy adult tissues. Snail1 reactivation is involved in fibrosis and in the progression of several cancer types^54^, as a potent driver of the EMT process in carcinoma cells^9^. Previous studies indicated that Snail1 induction in melanoma cells promotes invasion and metastasis^22^, however, Snail1 contribution to melanoma biology in an *in vivo* context was not defined. Our analyses of an inducible BRAF-driven melanoma reporter model reveal that SNAI1 in melanoma is reactivated in the stroma, particularly in CAFs. This is confirmed in syngeneic melanoma models where we find SNAI1 expression in CAFs in subcutaneous tumours and in lung metastases. In this study, we have generated mouse models that allow Snail1 ablation in an otherwise undisturbed immunocompetent environment to unveil the contribution of microenvironmental Snail1 to melanoma. We demonstrate that stromal Snail1 depletion blocks melanoma growth. This is associated with diminished proliferation and increased apoptosis of melanoma cells, pointing towards a non-cell autonomous role of microenvironmental Snail1 in melanoma cells. In accordance with this, Snail1 expression in CAFs from breast or colorectal cancer promotes epithelial cell invasion by paracrine signalling mediated by prostaglandinE2^11^. Here we show that Snail1-expressing CAFs mediate a tumour-promoting phenotype in melanoma by exerting an immunoregulatory role in the tumours.

Recently, single-cell sequencing technologies have shed light into the complexity and heterogeneity of CAFs in different tumour types and a better understanding of their functions and features can be harnessed to design better therapies for cancer treatments^16,50,55^. Although CAFs subsets in pancreas and breast have been characterised in detail^16^, less is known about CAFs modulation of melanoma biology. Recently, three different melanoma CAFs populations have been described^56^ and *Pdgfra* expression was widely found in the populations enriched at early stages of melanoma progression when our CAFs transcriptomic analysis was performed. Interestingly, we find that *Fap* is highly expressed in Snail1-expressing CAFs and downregulated upon its depletion. Further, we demonstrate that SNAI1 directly binds to the *Fap* promoter indicating that Snail1 can induce FAP expression in fibroblasts. Snail1 has classically been considered a potent transcriptional repressor^25^, however it can also act as a transcriptional activator^57,58^. In carcinoma cells, Snail1 directly activates the transcription of cytokines implicated in the recruitment of tumour-associated macrophages promoting TME remodelling^57^. Moreover, we previously showed that in renal fibrosis, Snail1 reactivation in tubular epithelial cells promotes a profibrotic inflammatory microenvironment by sustaining TGF-β signalling and cytokines production^52^. We show in this study that Snail1 depletion in CAFs also impinges on pathways associated to TGF-β signalling and inflammation and into the recruitment and activation of immune cells, indicating that Snail1 has a major immunoregulatory role when expressed in the melanoma microenvironment. Further, it is known that FAP-expressing CAFs populations are associated with immunosuppressive characteristics^47,48,50^, and that elevated FAP expression in CAFs through the JAK-STAT3 signalling pathway, contributes to a pro-tumorigenic immune response^45^. Consistently, our RNA sequencing data shows downregulation of the JAK-STAT3 signalling pathway among other pro-tumorigenic pathways when Snail1 is blocked. All these results are in line with the decrease in *Fap* levels we find in Snail1 KO-CAFs in association with anti-tumour immunity as shown by increased infiltration of cytotoxic CD8^+^-T, B cells and NK cells, and consistent with impaired melanoma growth. FAP^+^CAFs have also been associated with the recruitment of regulatory FOXP3^+^-T lymphocytes^48^, and in agreement with this, we find that the decrease in regulatory T cells we observe in the TME of tumours from Snail1^ME^-KO mice favours an anti-tumorigenic phenotype.

Interestingly, the effects of blocking Snail1 reactivation in the melanoma microenvironment are not restricted to the subcutaneous compartment but extend to the metastatic niche. Microenvironmental Snail1 depletion impairs the progression of experimental lung metastases associated with decreased *Fap* levels and anti-tumour immune responses. This suggests that despite CAFs heterogeneity, Snail1 is expressed in this population in an organ-independent manner to promote immunosuppression and tumour growth. We also demonstrate that *in vivo* systemic targeting with a Snail1 morpholino reduces metastatic burden in mice, extending mice survival. Moreover, as Snail1 was previously associated with increased metastases by favouring immune evasion by an EMT-dependent mechanism in melanoma cells^22^, our data further support Snail1 potential as a good therapeutic target in melanoma. Snail1 has been classically considered undruggable, however recently developed inhibitors of Snail1 protein-protein interactions have proved efficient in impairing tumour growth in mouse models of breast cancer^59^ and considering that Snail1 expression is almost absent in healthy tissues^54^, its inhibition in melanoma patients should be safe and lack major adverse effects.

The use of immune checkpoint inhibitors that target regulatory pathways on T cells to elicit antitumor responses has greatly improved the management of melanoma patients. However, only approximately 50% of patients respond^60^. Interestingly, Snail1-induced EMT in melanoma cells promoted resistance to immunotherapy based on intratumoural injection of dendritic cells^22^ and we show here that SNAI1 expression correlates with worse clinical responses to anti-PD-1 in melanoma patients. Current efforts directed to improve immune checkpoints inhibitors efficacy and the clinical management of patients include the characterization of mechanisms regulating immunosuppression^42^ and the discovery of biomarkers to predict responses. In this study, we show that Snail1 is a driver of CAFs-induced immunosuppression and pro-tumour immunity in melanoma, that its expression correlates with impaired responses to immune checkpoint inhibitors and therefore, we confirm its potential as a therapeutic target.

## Methods

### Mice

All experiments involving animals were performed in accordance with the European Community Council Directive (2010/63/EU) and Spanish legislation. The protocols were approved by the CSIC Ethical Committee and the Animal Welfare Committee at the Instituto de Neurociencias CSIC-UMH. Mice were hosted in a pathogen-free facility under controlled temperature, humidity, and 12 h light/dark cycle. All experiments were performed in 7-8-week-old mice C57BL/6. To analyse Snail1 in melanomas we crossed the inducible BRAF-driven mouse melanoma model BRafC^A^,Pten^loxP^,Tyr::CreERT2 (BRAF^V600E^/Pten^loxP^)^26^ (RRID:IMSR_JAX:013590) with Rosa-LSL-tdTomato (RRID:IMSR_JAX:007909) mice (referred as BRAF^V600E^/Pten^loxP^/tdTomato). To investigate Snail1 in the TME we crossed UBC-Cre-ERT2 mice^29^ (RRID:IMSR_JAX:008085) with Rosa-LSL-tdTomato (tdTomato) (RRID:IMSR_JAX:007909) and *Snai1^fl/fl^* mice^30^. To analyse myeloid populations in tumours, we used Cx3cr1CreERT2-YFP mice (RRID:IMSR_JAX:021160).

### Cell culture

Murine melanoma cell line BRAF^V600E^-5555^27,28^ were originally obtained from Richard Marais laboratory and luciferase-expressing Braf^V600E^-5555 (5555-Luc) were kindly given by Imanol Arozarena’s lab (NavarraBiomed). BRAF^WT^NRAS^WT^-B16F10 (CRL-6475) and BRAF^V600E^-YUMM1.7 (CRL-3362) cells were obtained from ATCC, and the FCT1 cell line was isolated from a tumour arising in BRAF^V600E^/Pten^loxP^/tdTomato transgenic mouse in our laboratory. NIH3T3 fibroblasts (CRL-1658) were purchased from ATCC. All cell lines were maintained in DMEM (Sigma) supplemented with 10% FBS (Sigma) and 1% penicillin/streptomycin (Sigma). Cells were kept at 37°C in a humid atmosphere containing 5% CO2 and the media was replaced every 2/3 days. Melanoma cells were passaged when they reached 80% confluency 1:10 every 72 h, while NIH3T3 cells were passaged when they reached 60-70% confluency 1:20 every 72 h. Cells were discarded up to seven consecutive passages and replaced by fresh stocks. All cell lines were tested and confirmed negative for mycoplasma monthly at the host institution.

### Inducible melanoma reporter model

Tumours were induced topically in 6-8 weeks BRAF^V600E^/Pten^loxP^/tdTomato mice. Treatment with 1.5μl 4 hydroxy tamoxifen (4-HT) (Sigma) (8 mg/ml), dissolved in ethanol:DMSO (80:20), was applied on the shaved skin of the back. Mice were immobilised until 4-HT dried completely. Tumours were collected when reaching approximately 1200 mm^3^ (formula: length x width x depth x 0.562).

### Melanoma subcutaneous allografts

5555 melanoma cells (5 x 10^6^ in 100ul in sterile PBS Ca^2+^Mg^2^-free) were subcutaneously injected in the dorsal area of Snail1^ME^-WT and Snail1^ME^-KO 7-8 weeks old mice. Treatment with tamoxifen (Sigma) (intraperitoneally, 100 mg/kg body weight), dissolved in corn oil:ethanol (90:10), was carried out to induce recombination. Tamoxifen administration began once tumours reached a volume of 80-100mm^3^. Tumour volume was recorded with a calliper every 2/3 days. When the tumours reached the limit size the mice were sacrificed, and tumours were collected for histological analysis.

For CAFs isolation by FACS Snail1^TME^-WT and Snail1^TME^-KO mice were injected with GFP-expressing 5555 cells as previously described. Four doses of tamoxifen were injected intraperitoneally on alternate days before collection and processing of the tumours. To study the myeloid populations in melanoma tumours, BRAF^V600E^-5555 melanoma cells were injected as described previously in Cx3cr1CreERT2-YFP mice.

### Experimental Metastasis Assay

To evaluate metastatic progression *in vivo,* Braf^V600E^-5555 -Luc (1 x 10^4^ cells in 100ul of sterile PBS Ca^2+^Mg^2^-free) were intravenously injected into the lateral tail vein, using a 27-gauge needle. Lung colonisation was analysed *in vivo* and *ex vivo* by BLI. Anaesthetized mice (isoflurane) were injected intraperitoneally with D-luciferin (Perkin Elmer) (150 mg/kg body weight) and imaged with an IVIS Lumina XR imaging system (PerkinElmer). The lung bioluminescence intensity signal of every mouse was determined using Living Image software (PerkinElmer). Tamoxifen treatment (intraperitoneally, 100 mg/kg body weight) was started once experimental metastases were established and detected by BLI imaging. Tamoxifen was administered three days a week until the end of the experiment. Mice were sacrificed after 3 weeks, and tissues were collected for histological analysis.

### Tumour processing

Tumours and lungs were fixed in 4% PFA for 4 h or ON respectively at 4°C. After fixation, tumours and lungs were washed three times with PBS and incubated in 30% sucrose for three days at 4°C before embedding in OCT. Embedded samples were kept in dry ice and transferred to −80°C before sectioning. Finally, OCT-embedded lungs and tumours were sectioned in a cryostat (Leica) at 8μm-thick sections and dried for 2 h at room temperature (RT) before being used for immunolabelling or stored at −80°C.

### Immunofluorescence (IF) stainings

Sections were blocked in 5% NGS, 1% BSA and 0.2% Triton x-100 for 1 h at RT and incubated with the primary antibodies O/N at 4°C in blocking solution and the following day for 30 min at RT. After extensive washing in PBS, slices were incubated with the secondary antibodies and DAPI in a blocking solution for 1h at RT. After washing the secondary antibody with PBS, slices were mounted in Dako Fluorescence Mounting Medium (Dako). Information and dilution of antibodies are listed in Table 1.

For IF in fibroblasts, FACs isolated cells were cultured and treated on poly-lysine-treated (Sigma) glass coverslips in 12-well plates and fixed with 4% PFA for 15 min at RT. Afterwards, cells were washed three times with PBS, permeabilized with 0.1% Triton x-100 in PBS for 15 min and blocked in a 0.1% Triton x-100 1% BSA solution for 1 h at RT. Then, cells were incubated with the primary antibodies O/N at 4°C in 1% BSA solution and the following day for 30 min at RT. After washing three times with PBS, cells were incubated with the secondary antibodies and DAPI 1 h at RT in 1% BSA solution. After washing the secondary antibody with PBS, cells were imaged.

Immunostainings were conducted using the primary and secondary antibodies listed in Table 1. Pictures were taken with an Olympus FV1200 confocal microscope with 20x or 40x objectives.

### Quantification of KI-67 and cleaved CASPASE-3

Proliferation and apoptosis were evaluated after IF staining by imaging sections and processing them with the ImageJ software. To analyse tumour proliferation and apoptosis, cell counts were obtained in 3 random fields from the tumour invasive front, 3 random fields from the tumour centre and 3 random fields from the tumour edge in each tumour slice. The same number of pictures were performed in every tumour slice. Four different tumours were analysed per condition. To analyse experimental lung metastases proliferation and apoptosis, representative pictures of different metastases were taken from each lung slice. Four different lungs were analysed per condition. The number of proliferating and apoptotic cells was determined as (number of apoptotic or proliferating tumour cells/ mm^2^).

### Histological analysis of Melanoma Lung Metastases

8um-thick lung sections were prepared and stained with Hematoxylin and Eosin (H&E) (Sigma) and documented with a Leica DFC700T digital camera. To quantify lung metastatic burden, 9 serial H&E stained lung sections were collected every 150um, spanning a total of 1200 microns of lung tissue. Total metastatic area (metastasis area/total lung area *100), number of metastases (number of metastases/ lung) and average metastasis size (total metastatic area/ number of metastases) were measured using Image J software.

### Tissue processing for flow cytometry

Tumours and lungs were mechanically disrupted using a scalpel blade followed by a cold and slow enzymatic digestion (2.5mg/ml Collagenase A and 0.2mg/ml DNAse I) (all from Roche) in PBS at 4°C for 1 h using constant gentle orbital agitation. After the incubation, the cell suspension was filtered through a 40um cell strainer using a 2ml syringe plunger. The content was centrifuged (5 min 350 g and 1 min 10.000 rpm) and pellets were resuspended in 1ml of RBC lysis buffer for 4 min at RT. Subsequently, cells were centrifuged and resuspended in fluorescence-activated cell sorting (FACS) buffer.

### FACS and flow cytometry

Prior to antibody staining, samples were blocked with Fc-block CD16/CD32 (Biolegend, 101320, 1:50) in FACS buffer for 10 min on ice to block nonspecific binding. For cell surface staining, cells were resuspended in the appropriate antibody cocktail and incubated for 30 min on ice protected from light. Samples were centrifuged and washed with a FACS buffer. For intracellular staining, cells were then collected and centrifuged for 5 min 350g. Cells were fixed, permeabilized and stained for transcription factors using the T rue-Nuclear Transcription Factor Buffer Set (Biolegend, Cat# 424401) according to the manufacturer’s instructions. Viability was assessed by staining with DAPI. Information and dilution for antibodies used for flow cytometry are listed in Table 1. For fibroblast sorting cells, PDGFRα^+^ GFP^-^ tdTomato^+^ were selected and sorted directly into a lysis solution from Arcturus PicoPure RNA Isolation Kit (Thermofisher). For sample validation, cells were plated in poly-L-lysine treated (Sigma) glass coverslips in 12-well plates and cultured for 24 h prior to IF. Immune cell profiling by flow cytometry was carried out by analysing 50.000 live singlets in each sample. All fluorescent data were analysed using BD FACSDiva Software (BD Bioscience).

### Total RNA extraction cDNA synthesis and qPCR analysis

RNA extraction from FACS-isolated samples was performed following the instructions in the Arcturus PicoPure RNA Isolation Kit (Thermo Fisher). The RNA was eluted in a volume of 15μl of elution buffer (TE) and 1μl was used for quantification and quality control using the Bioanalyzer High Sensitivity RNA chip. RNA extraction from bulk tumour or metastases samples was performed using the Illustra RNAspin Mini isolation kit (GE healthcare), following manufacturer’s instructions. For cDNA synthesis, Maxima First Strand cDNA Synthesis kit (Thermo Fisher) was used, following the manufacturer’s instructions. RT-qPCR was done using the Fast SYBR Green Mastermix (Applied Biosystems) and the primers listed in Supplementary Table 1. Relative levels of expression were calculated using a housekeeping gene and then experimental samples were normalised to their respective control.

### RNA sequencing

RNA degradation and purity were assessed using the RNA Nano 6000 Assay for the Bioanalyzer 2100 (Agilent). Samples were sent to Novogene Co. Sequencing libraries were generated using NEBNext® Single Cell/Low Input RNA Library Prep Kit for Illumina (NEB) following the manufacturer’s recommendations. Sequencing was performed using a cBot Cluster Generation Sequencing using PE Cluster Kit cBot-HS (Illumina) according to the manufacturer’s recommendation. After cluster generation, the library preparations were sequenced on an Illumina platform and 250 bp paired-end reads were generated.

### RNA sequencing Analysis

Raw data (raw reads) of FASTQ format were mapped to a mouse reference transcriptome (Mus_musculus.GRCm38.cdna.all.fa) built with Kallisto v.0.46.1. Read quantification to reference transcriptome was performed with Kallisto as well. The following steps were performed using R and RStudio. Tximport was used to import abundance.tsv files to R environment. EdgeR was used for differential expression analysis to obtain DEGlist objects and normalisation. The MatrixStats package was used to determine the statistics on the data. Data was filtered by choosing transcripts with at least 10 reads and later, at least 1 CPM in at least 3 samples. Normalisation factor TMM (trimmed mean of M-values) was applied. Limma and edgeR were used to obtain a final DEG list adjusted by BH (Benjamini-Hochberg) and sorted by p.value<0,05 and LFC >1. The graphical constructs of the RNAseq data were performed using gplots, plotly, gprofiler2, clusterprofiler and GSEABase.

### Plasmid constructs, interfering RNA and cell transfection

For RNA interference in NIH3T3 cells, siRNA obtained from Silencer® predesigned (Ambion) was used for Snail (Snail1 siRNA (antisense): AUAUUUGCAGUUGAAGAUCtt). SNAI1-Myc plasmid was transfected in NIH3T3 cells seeded in 6-well plates and 48 h after transfection cells were lysed for RNA extraction.

### Chromatin immunoprecipitation (ChiP) assay

NIH3T3 cells transfected with SNAI1-Myc were fixed at 80% confluency from 10 cm culture dish by adding 1% PFA for 10 min and subsequently quenched with glycine solution 0.125M for 5 min. Then, the cells were harvested and pooled together from 4 plates, and the chromatin was isolated using the Pierce™ Magnetic ChIP Kit (Thermo Fisher) following the manufacturer’s instructions. The sonication was performed in 15 cycles of 30-second on/off intervals in a Bioruptor® Pico sonication device (Diagenode). Finally, the immunoprecipitation and DNA isolation were performed using the same Pierce™ Magnetic ChIP Kit and the anti-Myc antibody listed in Supplementary Table 1. The isolated DNA was used for direct qPCR reaction.

### Vivo-Morpholino treatment

Snail1 Vivo-morpholino (Snail1-MO) (5’-TGAACTCTGCGGGAAGAGAAGAGAC-3’) against the boundary sequences of the intron 1 and exon 2 of Snail1 gene and standard control morpholino (Control-MO) that targets human β-globin intron mutation (5’-CCTCTTACCTCATTACAATTTATA-3’) were designed (by Gene Tools). C57BL/6 mice aged 7 weeks were injected in the tail vein with Braf^V600E^-5555 -Luciferase cells. 10 dpi, a solution containing Snail1-MO or Control-MO in saline (100 μl; 6 mg MO per kg) was injected in the tail vein of the corresponding mice every other day. After 20 days mice were sacrificed and lungs were processed, sectioned, and subjected to analysis.

### Statistical Analysis

All statistical tests were performed using GraphPad Prism 8 software. Student’s t-test or One-way ANOVA with Bonferroni’s multiple comparison test were performed to determine the significant values of the data. Kaplan-Meier data were analysed with the comparison of survival curves using the Long-rank (Mantel-Cox) test. All the values were shown as Mean values ± SEM (Standard Error of the Mean). Significant difference between groups were represented as follows: *= p≤0.05, ** = p≤0.01, *** = p≤0.001 and **** = p≤0.0001.

## Acknowledgments

We thank members of Berta Sanchez-Laorden and Angela Nieto’s labs for helpful discussions and comments along the project. We thank Joan Galceran and Khalil Kass Youssef for advice and help with the CHiP experiments. We thank the IN Imaging (Verona Villar and Giovanna Exposito) and OMICS (Antonio Caler) facilities for the technical support. This work was supported by grants SAF2016-75702-R and PID2019-106852-RBI00 (to B.S-L), RTI2018-096501-B-I00 and PID2021-125682NB-I00 (to M.A.N) funded by MCIN/AEI/ 10.13039/501100011033 and ERDF, the Melanoma Research Alliance (https://doi.org/10.48050/pc.gr.91574 to B.S-L), the FERO Foundation (to B.S-L) and the AECC Scientific Foundation (FC_AECC PROYE19073NIE to M.A.N). M.A-P and F.C-T were recipient of a FPI predoctoral contract. FJR-B and F.G-Q hold contracts from the Generalitat Valenciana (APOSTD and ACIF respectively). We also acknowledge financial support from the Spanish State Research Agency, through the “Severo Ochoa Program” for Centres of Excellence in R&D (SEV-2017-0273).

## Author contributions

M.A-P. performed most experiments, analysed, and interpreted the data, and contributed to writing the manuscript. F.J.R-B. analysed and interpreted the transcriptomic data and performed some experiments, F.C-T., F.G-Q and C.L-B performed experiments. M.A.N provided the UBQ-Cre-Snail1^loxp^ mice and helped in the interpretation of data. B.S-L. conceived the project, interpreted the data, and wrote the manuscript.

## Data availability statement

Datasets related to this article will be made available via a public repository upon acceptance of the manuscript or as required by the editorial office.

---

**Supplementary Figure 1 (Related to Figure 1).**
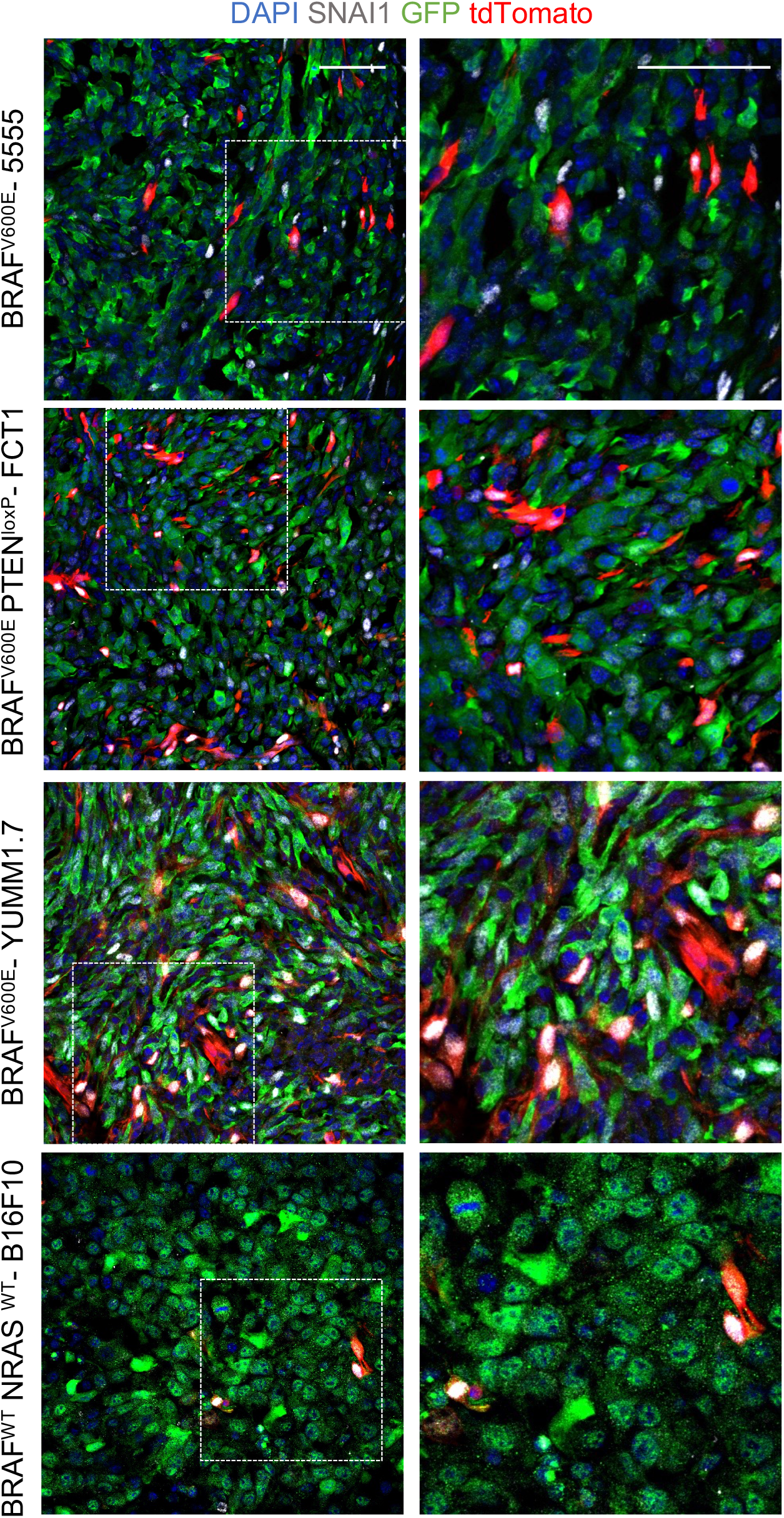
Snail1 expression in different syngeneic melanoma models. Representative images of double immunolabeling for SNAI1 (white) and GFP (green, melanoma cells) in subcutaneous tumours in tamoxifen treated Snail1^ME^-WT (stromal cells in red) upon injection of different mouse melanoma cell lines tagged with GFP: 5555 (BRAF^V600E^); FCT1 (BRAF^V600E^PTEN^flox/+^); YUMM1.7 (BRAF^V600E^PTEN^flox/flox^Cdkn2a^-/-^); B16F10 (BRAF^WT^NRAS^WT^). Scale bars: 50μm

**Supplementary Figure 2 (Related to Figure 2).**
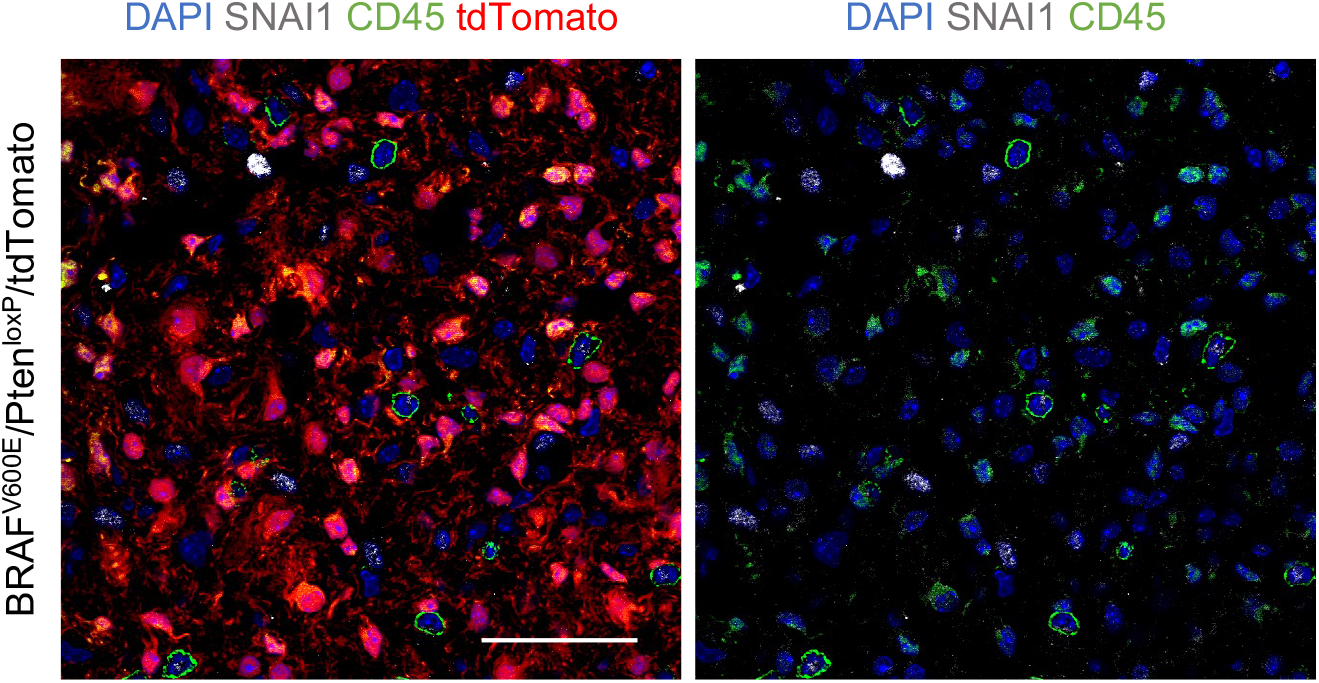
Snail1 is absent in immune cells in tumours from an inducible BRAF^V600E^/Ptenl^oxP^/tdTomato melanoma model. Representative images of double immunolabeling for SNAI1 (white) and CD45 (green, immune cells). TdTomato indicates melanoma cells (red). Scale bar: 50μm.

**Supplementary Figure 3 (Related to Figure 2).**
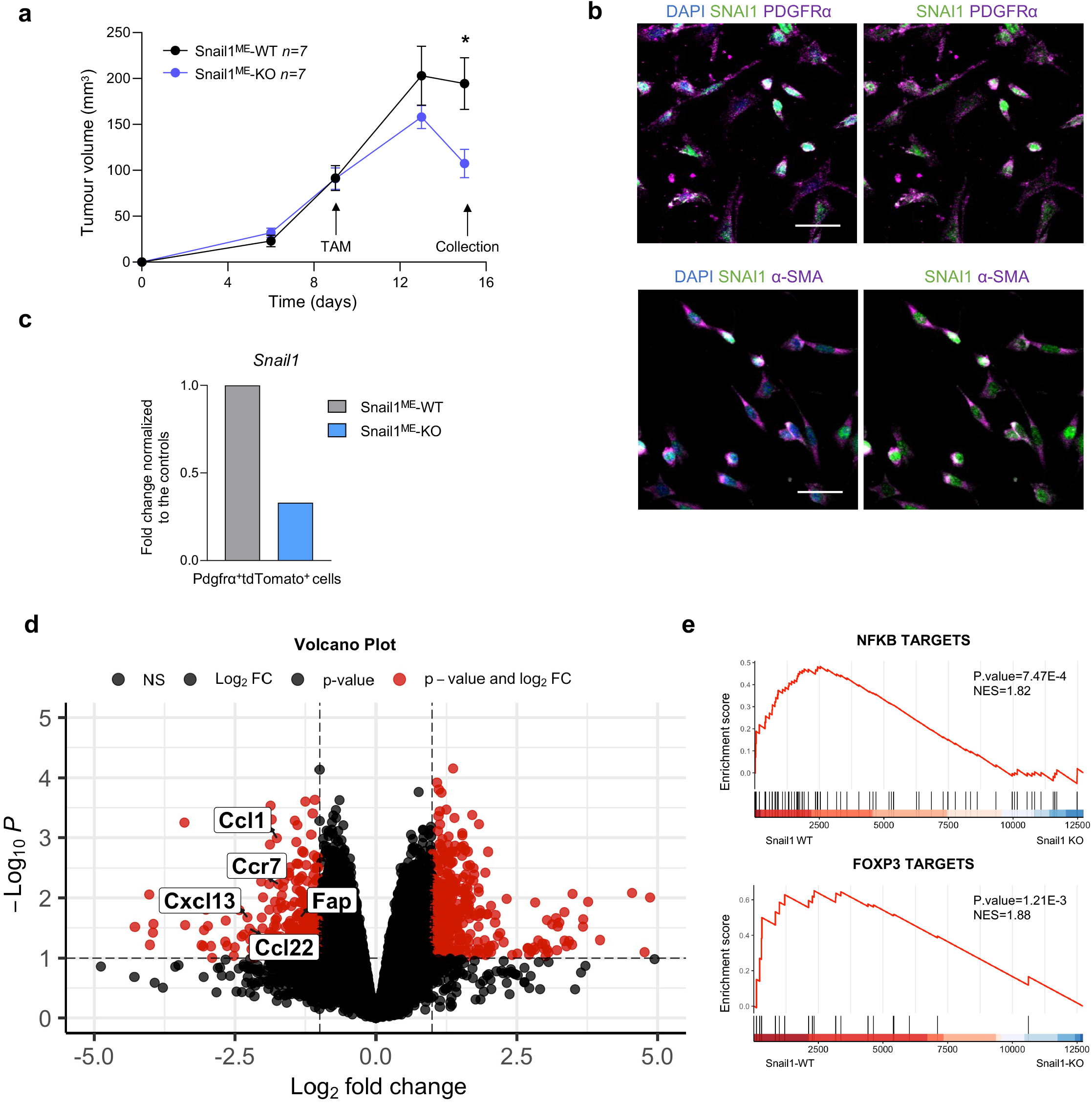
Isolation strategy and transcriptomic analysis of Pdgfrα^+^ CAFs from Snail1^ME^-WT and Snail1^ME^-KO mice. **(a)** In vivo experiment designed to isolate fibroblast from Braf^V600E^-5555 melanomas grown subcutaneously in Snail1^ME^-WT and Snail1^ME^-KO mice. Tumour growth graph of an experiment showing the selected time point for the CAFs isolation (Snail1^ME^-WT=7 and Snail1^ME^-KO=7). Data are represented by Mean±SEM and statistically significant differences are tested by unpaired two-tailed Student t-test (*=p<0.05). **(b)** Representative images of double immunolabeling for SNAI1 (green) and different CAFs markers (magenta), in cells isolated from tumours in (a) after FACS-sorting (Pdgfrα^+^tdTomato^+^GFP^-^). Scale bars: 50μm. **(c)** *Snail1* mRNA levels assessed by RT-qPCR to validate the fibroblasts population isolated by FACS (Pdgfrα^+^tdTomato^+^GFP^-^) from tumours in (a) (samples from 3 animals with the same genotype were pooled for each condition). **(d)** Volcano plot of Log_2_ fold change of DEGs between Snail1-WT and Snail1-KO fibroblast samples. The red dots on the right represent the upregulated genes and the red dots on the left the downregulated genes. **(e)** Gene set enrichment analysis (GSEA) showing enrichment of the indicated signatures in the Snail1^ME^-WT and Snail1^ME^-KO CAFs from tumours. NES, normalised enrichment score.

**Supplementary Figure 4 (Related to Figure 5).**
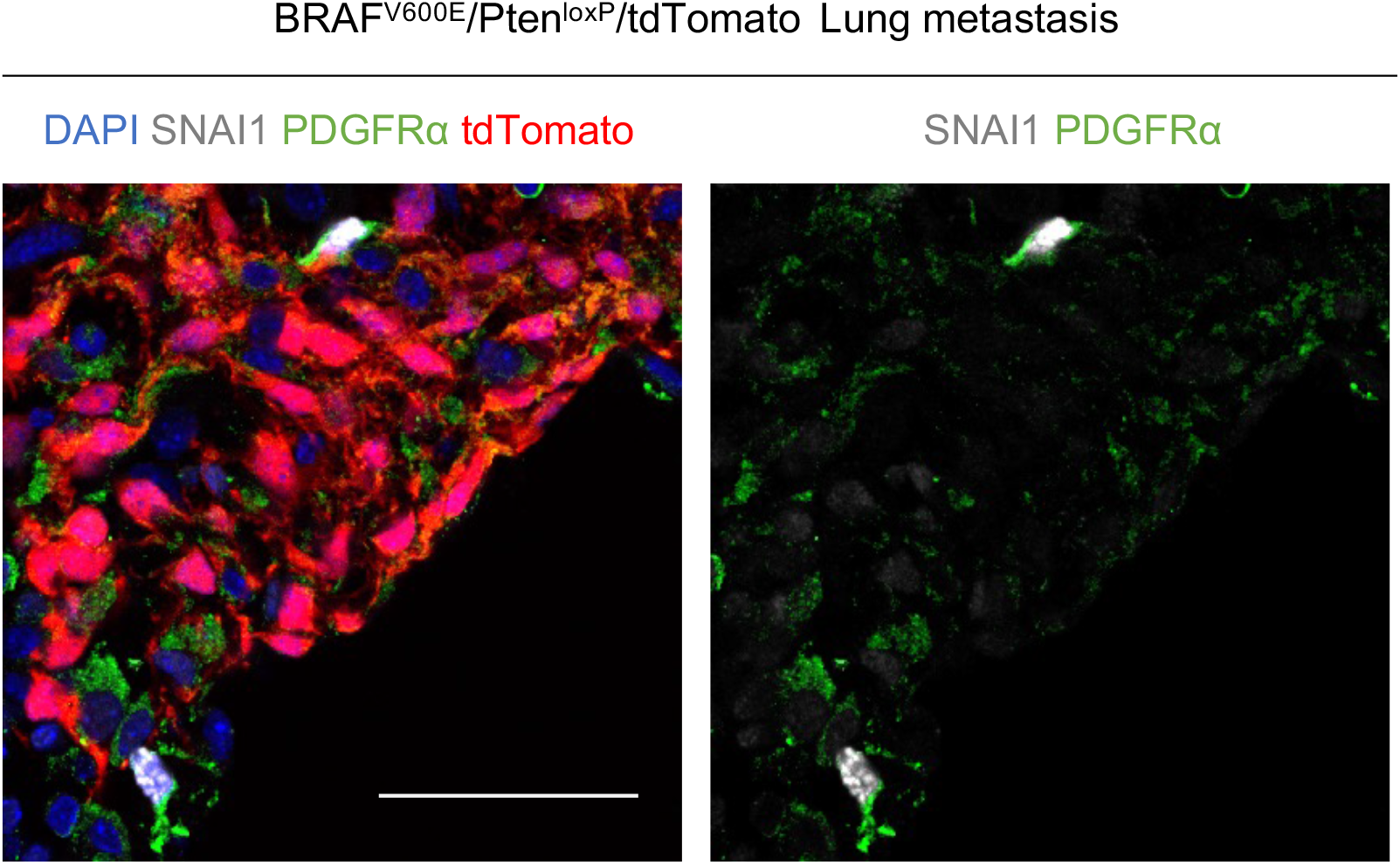
Snail1 is expressed in Pdgfrα^+^CAFs in tumours from an inducible BRAF^V600E^/Ptenl^oxP^/tdTomato melanoma model. Representative images of double immunolabeling for SNAI1 (white) and PDGFRα (green, fibroblasts). tdTomato indicates melanoma cells (red). Scale bar: 50μm.

**Supplementary Table 1.**
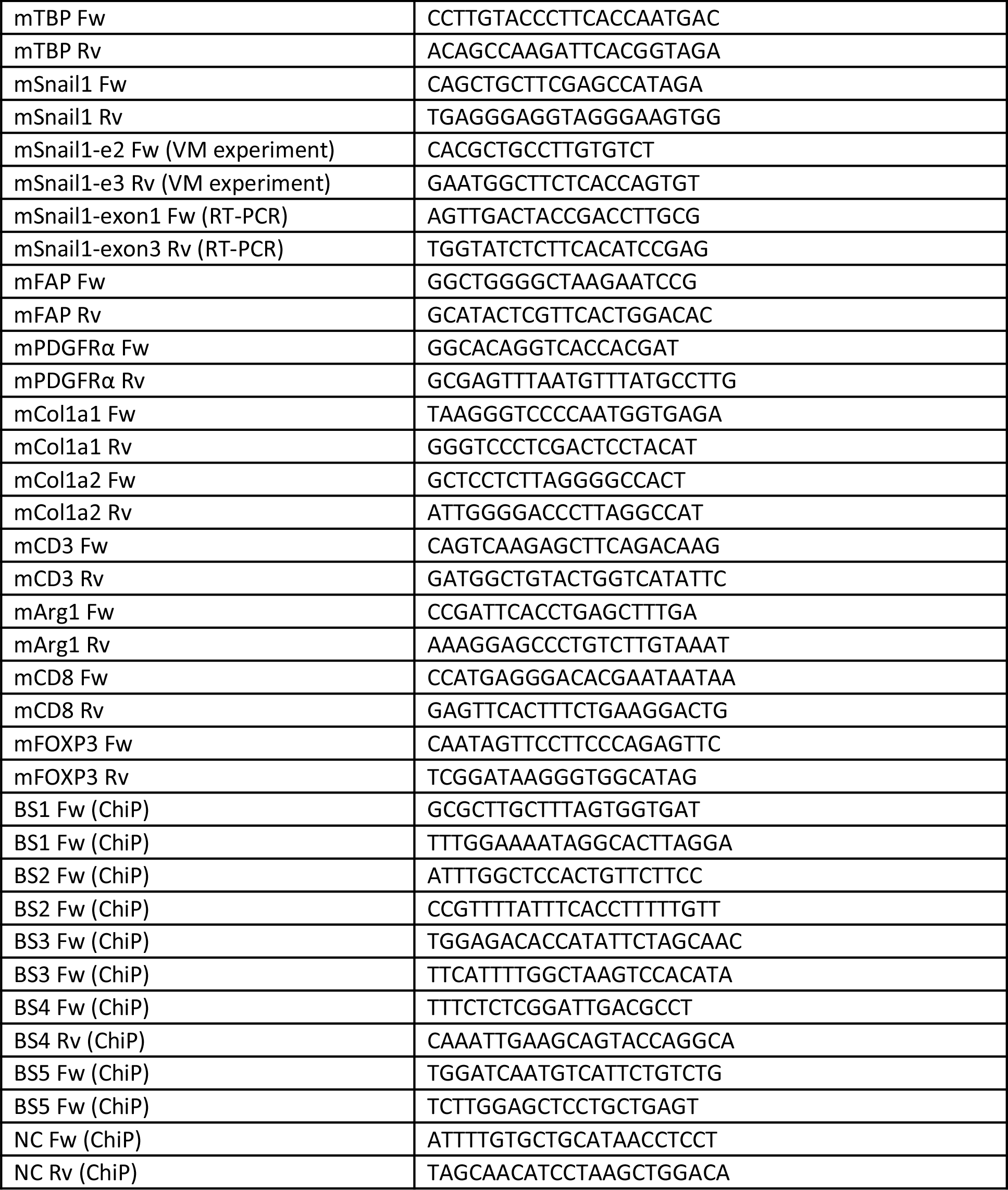
Primer sequences

